# Spatially structured competition and cooperation alters algal carbon flow to bacteria

**DOI:** 10.1101/2024.06.14.598523

**Authors:** Hyungseok Kim, Vanessa L. Brisson, John R. Casey, Courtney Swink, Kristina A. Rolison, Amber N. Golini, Trent R. Northen, Peter K. Weber, Dušan Veličković, Cullen R. Buie, Xavier Mayali, Rhona K. Stuart

## Abstract

Microbial communities regulate the transformations of carbon in aquatic systems through metabolic interactions and food-web dynamics that can alter the balance of photosynthesis and respiration. Direct competition for resources is thought to drive microbial community assembly in algal systems, but other interaction modes that may shape communities are more challenging to isolate. Through untargeted metabolomics and metabolic modeling, we predicted the degree of resource competition between bacterial pairs when growing on model diatom *Phaeodactylum tricornutum-*derived substrates. In a subsequent sequential media experiment, we found that pairwise interactions were consistently more cooperative than predicted based on resource competition alone, indicating an unexpected role for cooperation in algal carbon processing. To link this directly to algal carbon fate, we chose a representative cooperative and competitive ‘influencer’ isolate and a model ‘recipient’ and applied single-cell isotope tracing in a custom porous microplate cultivation system. In the presence of live algae, the recipient drew down more algal carbon in the presence of the cooperative influencer compared to the competitive influencer, supporting the sequential experiment results. We also found that total carbon assimilation into bacterial biomass, integrated over influencer and recipient, was significantly higher for the cooperative interaction. Our findings support the notion that non-competitive interactions are critical for predicting algal carbon fate.

**Significance Statement:** Microbial interactions have widely been studied in the context of host resources but testing and measuring direct interactions in a lab has been particularly challenging. By combining untargeted metabolomics, sequential/(co-)culture, and metabolic modeling, we demonstrate that the presence of an unexpected interaction mode in a live system and show how it impacts the flow of host-derived resources. This top-down approach can help identify novel bacterial interactions that play a crucial role in microbial community-host ecosystems, which may have an impact in holobiont phenotypes including alga, fungal, or plant systems.

## Introduction

Most algal-associated bacteria subsist on microalgal exudates, but how bacteria-bacteria interactions influence algal carbon (“C”) fate is not well constrained. Microbial interactions can have profound implications for carbon flux in diverse ecosystems. For example, one recent study found that microbial respiration is dependent on the degree of cooperation and competition among bacteria (1). When examining algal carbon derived food webs, a consumer-resource based model is typically used as the basis for prediction. Indeed, host resources alone have been shown to be predictive of algal-associated microbial community assembly (2). At the base of the food web, (algal C directly to bacterial taxa), host resources could be partitioned based on resource consumption ability, also known as exometabolic niche partitioning (3), which divides bacterial taxa into functional guilds. However, these functional guilds alone do not always predict resource consumption outcomes, and incorporation of bacteria-bacteria interactions into models can improve disparities in predictions based solely on consumer-resource models (4).

Bacteria-bacteria interactions can be broadly grouped into competitive and cooperative forms, with a variety of mechanisms underlying each type. Competition for host resources (referred to herein as ‘resource competition’) can be predicted by degree of metabolic niche partitioning and then tested with pairwise interaction experiments (e.g. (5)). Thus, resource competition is generally considered the most straightforward of bacterial interactions to measure and compare (6), and bacterial metabolic niche partitioning has been well characterized in regards to algal dissolved organic carbon consumption (7, 8). Interference competition involves direct cell damage between individuals with a variety of mechanisms (9), and is also considered to be widespread (10, 11). Although cooperative metabolic interactions are more difficult to detect, metabolic cross feeding is thought to be ubiquitous (12), and there are examples of syntrophic breakdown of complex organic matter (13-17). Cooperative bacterial interactions can also include production of facilitator exometabolites (e.g. siderophores (18-20)), or breakdown/uptake of toxic compounds (21-23)). Along with experimental evidence of these interactions, genome-based predictions of metabolic overlap has also been broadly used to predict resource competitive and as well as metabolic cross-feeding based cooperative interactions (17, 24-28). Thus, while consumer-resource based predictions, and resource competition in particular, could be considered a baseline, the influence of other interaction types are less well understood.

Here, we sought to quantify cooperative bacteria-bacteria interactions in a model algal community, in a mechanism-independent manner, and then compare algal C fate of a cooperative interaction to a baseline resource competitive interaction. We used a set of bacterial algal isolates that are all able to grow with the diatom *P. tricornutum* without any exogenous organic carbon added (29). In earlier work in an enrichment community containing these isolates growing with *P. tricornutum*, we found that while exogenous resource additions could promote some taxa based on their consumption abilities, other taxa that could not consume the resource also were promoted. In addition, some taxa that could consume the resource were excluded, indicating the importance of both cross-feeding and resource competition in community structuring (30). Since the diatom and its exudates are the sole source of organic C in this system, we set an assumption of a resource-consumption based model of community assembly, with resource competition between bacterial taxa as the baseline for all bacteria-bacteria interactions. We then used metabolomic profiling and genome-based metabolic models to predict the degree of potential resource competition between all bacterial pairs, which we then tested in a sequential spent media experiment. The outliers from our predictions thus represented either cooperative interaction or interference competition, based on which way they deviated from our predictions, giving us a degree of potential cooperation between bacteria when grown with algal exudate. To quantify the role of cooperative interactions in algal C fate, we took the most cooperative pair and the pair that best matched our resource competition prediction, and quantified algal exudate consumption *in vivo* between the different partners, using a custom porous microplate system (31-33) and stable isotope probing paired with high-resolution imaging mass spectrometry (34-37).

## Results

### Metabolite consumption patterns and metabolic models predict degree of resource competition

Bacteria grown on *P. tricornutum* spent medium consumed a subset of algal exometabolites. Our analysis detected 162 LC-MS/MS features (hereafter referred to as metabolites), based on retention time and m/z values, that were significantly (Bonferroni adjusted P-value < 0.05 from Student’s t-test) above background compared to our extraction blanks for at least one sample group (bacterial isolate or *P. tricornutum* spent medium) (Supplementary Table S1, Supplementary Figure S1). We note that the extraction and analytical methods used here lead to poor detection of small, highly polar compounds and thus our data represent a subset of all possible exometabolites (38). A subset consisting of 54 (33%) metabolites were significantly changed (either increased or decreased in relative signal intensity) for at least one bacterial isolate compared to uninoculated *P. tricornutum* spent medium (Figure 2a). Of these 54, seven (13%) could be putatively identified based on their MS/MS spectra (Table 1, with identification details in Supplementary Table S2). Bacterial isolates produced and consumed distinct suites of metabolites (Figure 2a). Only four metabolites were depleted by at least half (log_2_ fold change ≤ - 1) by all bacterial isolates, and no metabolites were universally produced by all isolates (Figure 2a). Notably, two of the four metabolites consumed by all isolates were putatively identified as dipeptides. Hierarchical clustering grouped the bacterial isolates based on their patterns of production and consumption (Figure 2a, left dendrogram). A Mantel test (39) was conducted to assess whether the patterns of metabolite consumption correlated with bacterial phylogeny, and no phylogenetic correlation was detected (r = 0.11, *p*-value = 0.29).

**Table 1.** Putative identifications (based on MS/MS spectral matches to GNPS) of consumed or produced metabolites.

| Metabolite Feature ID | Putative Identification | Consuming strains | Producing strains |
| --- | --- | --- | --- |
| Positive-1492 | Val-Leu or Val-Ile | All isolates |  |
| Positive-7891 | Val-Phe | All isolates |  |
| Positive-21 | Thymine | All isolates except *Marinobacter* 3-2, *Labrenzia* 13C1 and *Oceanicaulis* 13A |  |
| Positive-31 | Guanosine | All isolates except *Marinobacter* 3-2, *Labrenzia* 13C1 and *Oceanicaulis* 13A | *Marinobacter* 3-2 and *Oceanicaulis* 13A |
| Positive-37 | Phenylalanine | All isolates except *Devosia* B7WZ |  |
| Negative-20 | Deoxyguanosine |  | *Marinobacter* 3-2 and *Labrenzia* 13C1 |
| Positive-23 | Absciscic acid |  | *Alcanivorax* EA2 |

**Figure 1.**
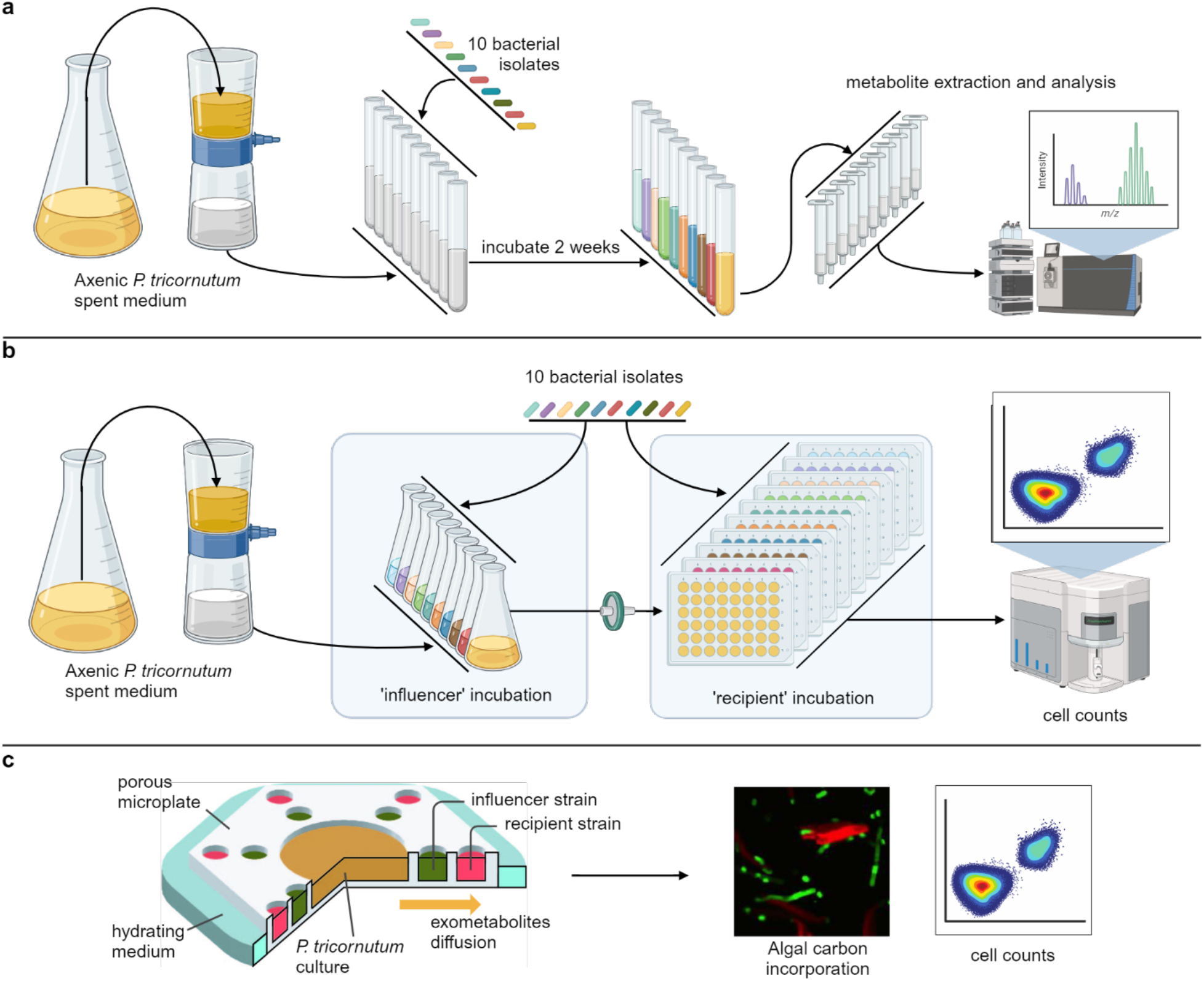
Testing bacterial-bacterial interactions with diatom *P. tricornutum*. Experimental procedures to (a) profile differential metabolite consumption and production by each bacterial isolate using untargeted metabolomics, (b) quantify isolate abundances through spent medium exchange, and (c) measure isolate activity and bacteria-bacteria interaction in response to *in vivo* algal exometabolites in porous microplates.

**Figure 2.**
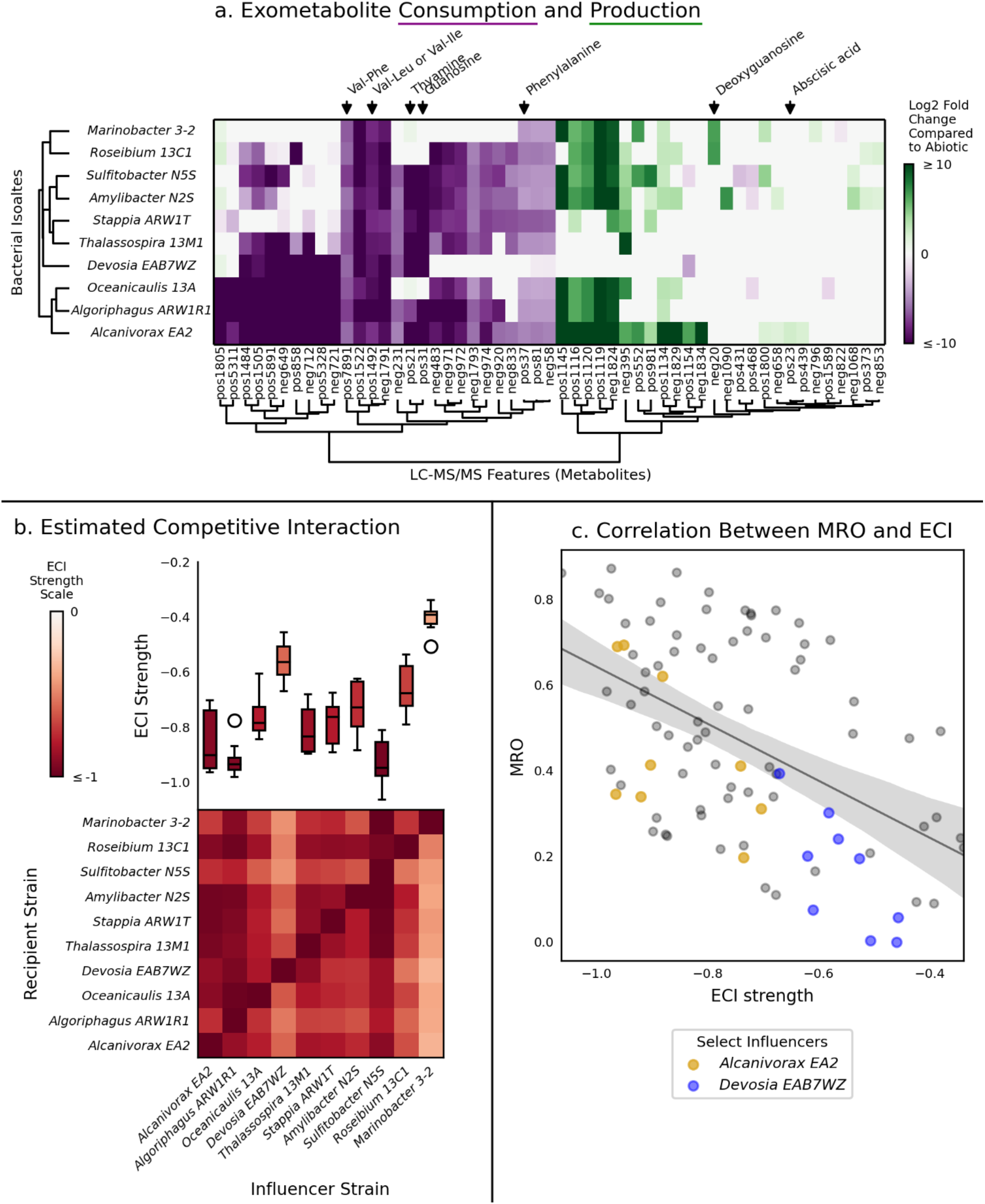
Bacterial resource competition for algal exometabolites. (a) Metabolite consumption and production by bacterial isolates. Columns represent each metabolite, with putative identifications (based on MS/MS spectral matches to GNPS) indicated at the top. Rows represent each bacterial isolate inoculated into *P. tricornutum*-spent medium. Colors represent statistically significant (*P* ≤ 0.05 for Student’s *t*-test with Bonferroni correction) consumption (purple) or production (green) of each metabolite compared to the levels in uninoculated *P. tricornutum* spent medium. Color lightness/darkness indicates the extent of consumption or production (log2 fold change compared to uninoculated *P. tricornutum* spent medium), with darker colors indicating greater changes. White indicates no significant difference. Left dendrogram shows hierarchical clustering of bacterial isolates based on metabolite feature consumption and production profiles. Bottom dendrogram shows hierarchical clustering of LC-MS/MS features (b) Estimated competitive interactions (ECI) based on metabolite consumption overlap. Heatmap shows the ECI for each influencer strain (columns) on each recipient strain (rows). Darker red indicates stronger competitive interactions. Top boxplot summarizes ECI for each influencer strain (column). Boxplot colors indicate median ECI values. (c) Correlation between metabolic resource overlap (MRO) and ECI. Each point represents one influencer-recipient pair. Influencers studied in the porous microplate experiment are highlighted in yellow and blue for *Alcanivorax* EA2 and *Devosia* EAB7WZ respectively. The diagonal line shows the Pearson correlation (R^2^ = 0.25, *P* = 6 × 10^-7^), and the grey area indicates the 95% confidence interval. Degree of resource competition is predicted by the metabolite patterns and the metabolic models.

We characterized an overlap between metabolite consumption patterns by calculating a predicted competitive interaction strength (ECI) between influencer and recipient strains (5, 40, 41), where the influencer has access to the algal metabolites before the recipient, and found a wide range of resource competition (Figure 2b). For instance, *Marinobacter* 3-2 and *Devosia* B7WZ as the influencer strains, had the weakest (least negative) ECI strengths suggesting they are less likely to compete with others. This is in contrast with three other strains (*Alcanivorax* EA2, *Algoriphagus* ARW1R1, and *Sulfitobacter* N5S) as influencer where they showed stronger competitive interaction strengths with other strains. As recipient strains, on the other hand, most isolates had a highly variable ECI, depending on the influencer strain.

Since all of these strains have genomes available, we independently calculated the metabolic resource overlap (MRO), a modeling-based estimate of how much two organisms can use the same resources, and found agreement with the ECI-based estimate (Figure 2c, Supplementary Figure S2). The MRO and ECI were significantly negatively correlated (Pearson’s R^2^ = 0.25, P = 6 × 10^-7^), indicating that a modest portion of the observed competitive interactions could be predicted based on known metabolic potential alone.

### Sequential bacterial interactions were more positive than predicted from resource competition

After obtaining a baseline estimate of resource competition among the algal-associated bacteria (based on both ECI and MRO scores), we conducted a sequential spent medium experiment to both test these predictions and look for outliers that might indicate the presence of other bacteria-bacteria interactions. This experiment tested whether and to what extent growth of one bacterial strain on *P. tricornututm* spent medium and consumption of metabolites in the first phase of the experiment would negatively impact the growth of a second bacterial strain in the second stage of the experiment. We calculated a sequential interaction strength (SI) from the cell counts of the second strain (Figure 3a, Supplementary Table S3) (5).

**Figure 3.**
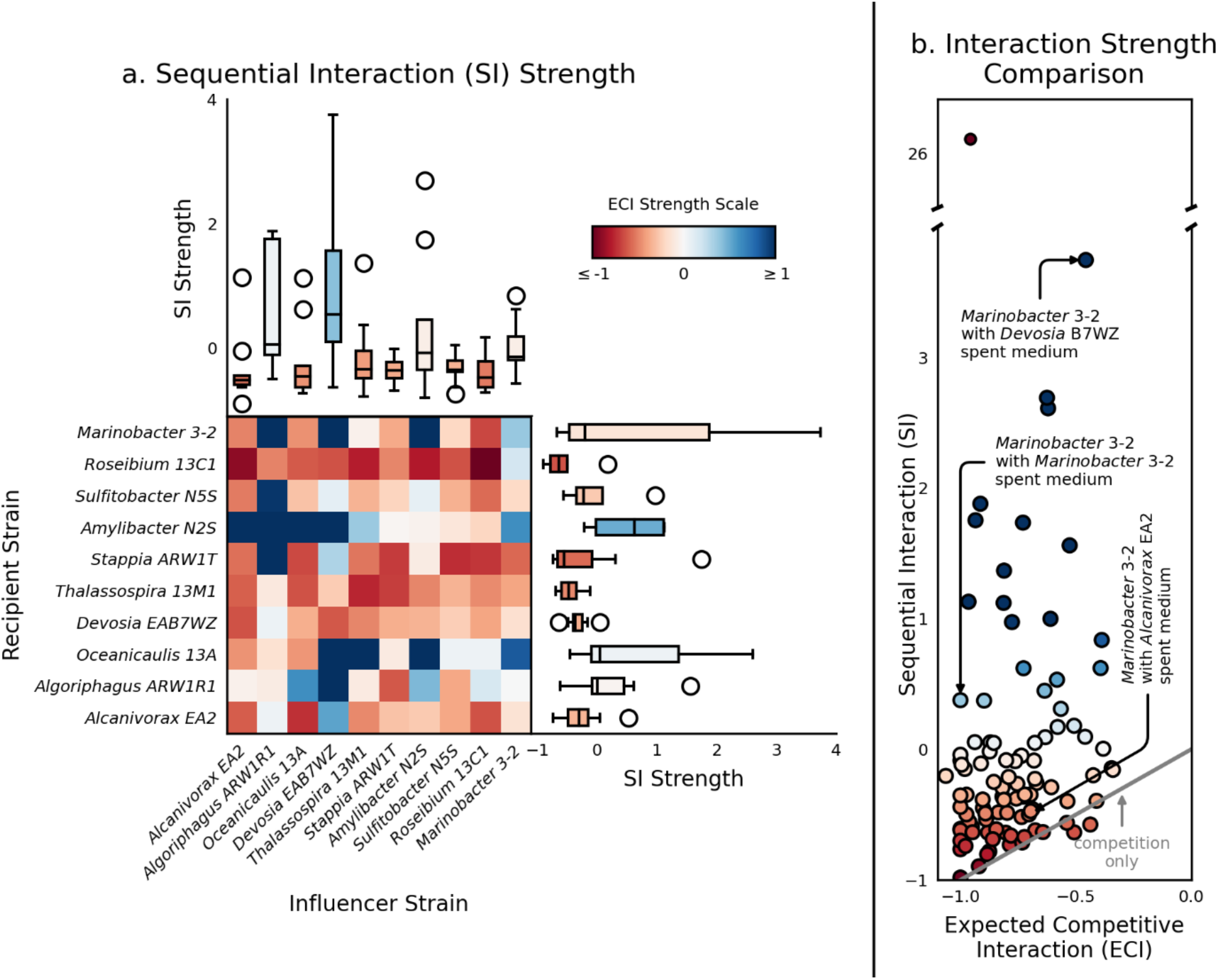
Sequential interactions (SI) in comparison to expected competitive interactions (ECI). (a) SI based on growth from sequential spent medium exchange. Heatmap shows the SI for each influencer strain (columns) on each recipient strain (rows). Darker red indicates stronger negative interactions. Darker blue indicates stronger positive interactions. Top boxplot summarizes SI for each influencer strain (column). Right boxplot summarizes SI for each recipient strain (row). Boxplot colors indicate median SI values. (b) Comparison of measured sequential interaction (SI) strengths to expected competitive interaction (ECI) strengths. Each point represents the SI and ECI for one bacterial influencer/recipient pair. Point coloring indicates the strength of a negative (red) or positive (blue) SI. The diagonal line indicates the expected competition only case where SI = ECI. Annotations indicate bacterial isolate pairs selected for porous microplate experiments. Sequential spent medium exchange suggests the presence of outliers and other bacterial-bacterial interactions.

We found some of the observed patterns in SI strengths were consistent with the ECI strengths based on metabolite consumption (Figure 3a). For instance, as with the ECI, *Devosia* B7WZ and *Marinobacter* 3-2 had generally high (more positive) SI strengths as influencers, while *Alcanivorax* EA2 had consistently low SI strengths, indicating generally negative influences on most other isolates.

However, although competition was evident in some sequential interactions, the results also indicated the importance of other factors contributing to less negative interactions than predicted from resource competition alone (Figure 3b). A comparison of ECI and SI for all pairwise combinations of influencer and recipient strains showed that while some SI matched closely with the ECI, falling along the 1:1 diagonal line, most were above the line, indicating that the SIs were less negative than expected from the ECIs. In fact, some SIs were highly positive, indicating growth promotion of the second stage bacterial isolate by growth in spent medium from the first stage isolate. Reflecting this variation, we did not observe an overall correlation between SI and ECI (p = 0.8).

### Designing custom porous microplate experiments to test bacterial interactions in situ

Our sequential interaction experiment indicated the potential prevalence of cooperative bacteria-bacteria interactions for incorporation of algal organic matter, on top of the predicted resource competition, so the next experiment set out to both confirm this interaction in the presence of live algae and quantify and compare algal C fate relative to resource competition only. We chose the most cooperative influencer, *Devosia* B7WZ as a model, and chose *Alcanivorax* EA2, which exhibited the least cooperative interaction (based on average influencer SI strength, Figure 3a). *Alcanivorax* EA2 was also chosen because its sequential interactions best matched predictions based on ECI and MRO, and it consumed the most diverse set of metabolites among the isolates, suggesting the resource competition alone likely drives its interactions. *Marinobacter* was chosen as the model recipient as it had the broadest range of interaction values (Fig. 3b), indicating it is sensitive to influence (Figure 4a). We also included a positive control with *Marinobacter* as both the influencer and recipient (maximum resource competition), and a negative control with no influencer (Figure 4a). We then designed a custom porous microplate, which allows the exchange of nutrients or metabolites while physically separating the isolates and algae (32). To ensure sequential bacterial uptake of the algal exometabolites in the porous microplate, a parameter study was conducted using a back-of-the-envelope calculation of algal DOC reaching to the surrounding wells in the device. Based on previous measurements of microbial growth (32, 33) and biomolecular diffusion in the copolymer (31), we estimated the DOC concentration under various geometrical configurations of a microplate (Supplementary Figure S3, Supplementary Note S1). The hexagonal array of wells with their center-to-surrounding ratio ∼3:1 was designed so that algal C flux to bacteria is like *in situ* algal-bacterial co-culture. The design was used to fabricate the culture devices made of nanoporous copolymer poly(2-hydroxethyl methacrylate-*co*-ethylene glycol dimethacrylate) (see methods) (31-33). We then incubated the host *P. tricornutum* and two bacterial isolates in the custom porous microplates (Figure 4a). To track photosynthetically-fixed carbon and examine metabolic activities of the isolates, ^13^C-labeled bicarbonate and ^15^N-labeled leucine were added to the incubations. All influencers in the inner wells incorporated a higher amount of algal C at the single cell than recipients in the outer wells (Supplementary Figure S4), supporting our microplate design configuration.

**Figure 4.**
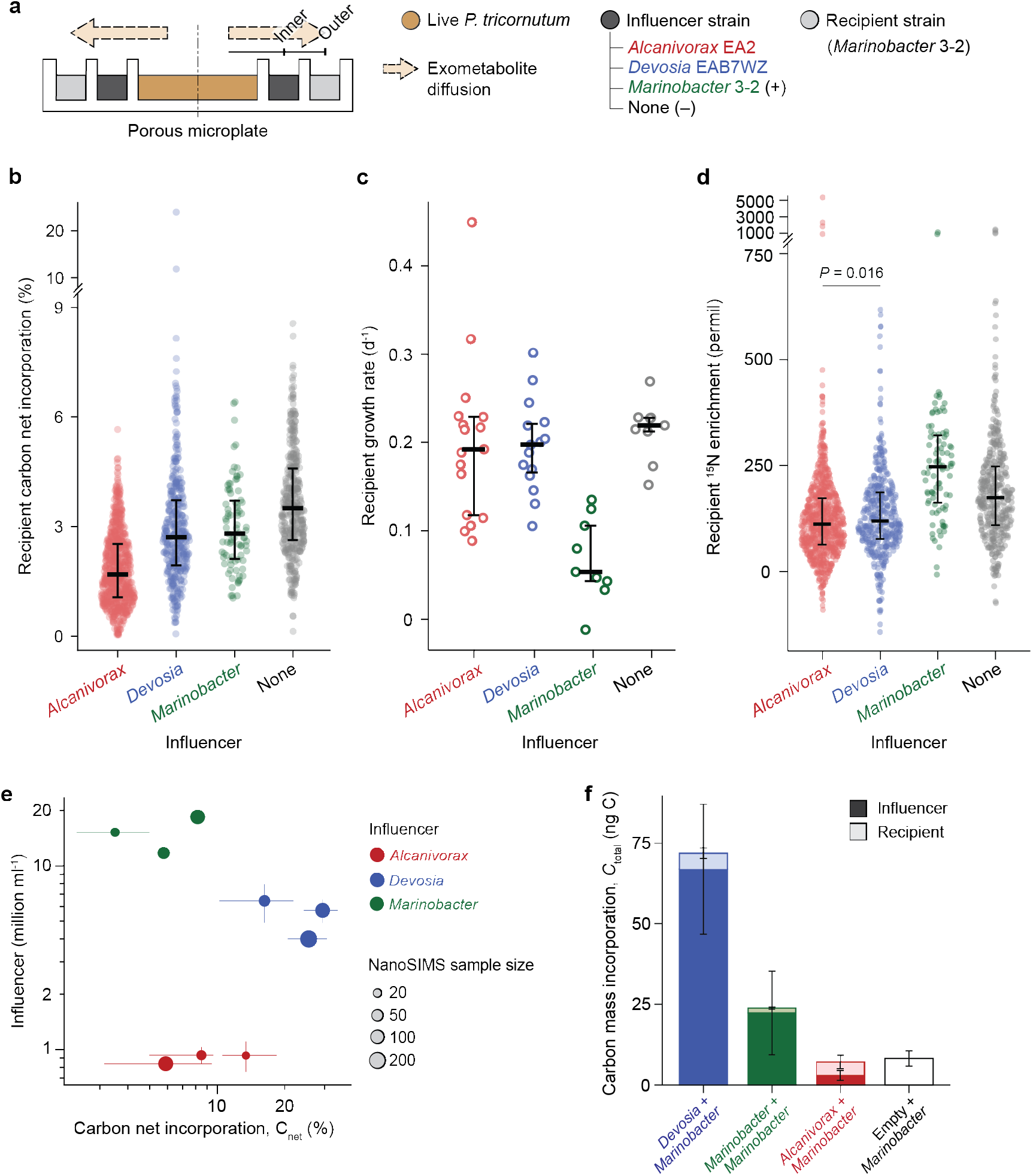
Porous microplate to test interspecies interactions. (a) Side view schematic of porous microplate for co-culturing *P. tricornutum* and bacterial influencer/recipient. The microplate was designed in a way that algal C flux to bacteria is like in situ algal-bacterial co-culture by maximizing center-surrounding well volume ratio. (b) Net carbon incorporation (C_net_) by recipient strain *Marinobacter* on day 14. Each point represents a single cell NanoSIP measurement. (c) Average growth rate of the recipient *Marinobacter* from day 5 to 14, co-cultured with influencer strains and *P. tricornutum* in the porous microplate. Each point represents a measurement from microplate well. (d) Enrichment of ^15^N by recipient strain *Marinobacter* on day 14. Each point represents a single cell NanoSIP measurement. Black lines indicate median (thick) and interquartile range (error bar). (e) Relationship between bacterial growth rate and single cell C_net_ for influencer strains after 14 days. Each symbol represents measurement from a single microplate. Vertical and horizontal error bars respectively denote standard deviation of cell number and the first/third quartile of C_net_. (f) Estimate of total incorporation of the carbon mass (*C*_total_) by bacterial pair (see Supplementary Note S2 for derivation). Error bars denote standard deviation of three microplate replicates. The microplate co-culture confirms the presence of bacterial interactions with live *P. tricornutum*.

### Recipient strain incorporated more algal carbon with cooperative influencer than competitive influencer

In our porous microplates, we found that the recipient *Marinobacter* assimilated more algal-derived carbon with influencer *Devosia* in the inner ring relative to influencer *Alcanivorax* (Figure 4b). From single cell ^13^C enrichment values, we calculated the percent of bacterial carbon derived from the algal C, C_net_ (29, 35, 42). Median C_net_ for *Marinobacter* when *Devosia* was the influencer was significantly higher (2.71%) than when *Alcanivorax* was the influencer (1.69%, *P* < 0.0001, Wilcoxon test), supporting a cooperative interaction between *Devosia* and *Marinobacter* (Figure 4a). However, from the negative control, we found that the presence of any influencer lowered the C incorporation of *Marinobacter* cells (*P* < 0.001, Kruskal-Wallis test). This supports a role for resource competition as underlying most interactions, suggesting that even our cooperative influencer removes some carbon substrates that *Marinobacter* would otherwise consume. From bacterial abundances in the wells over the incubation, we calculated bacterial growth rates, but did not observe a significant effect on *Marinobacter* growth in the presence of *Devosia* or *Alcanivorax* relative to the negative control (*P* = 0.511, 0.358, Wilcoxon test, Figure 4c). However, the positive control with *Marinobacter* in the inner well did exhibit a significantly reduced growth rate (0.054 d^-1^, *P* < 0.001, Wilcoxon test, Figure 4c, Supplementary Figure S5), as expected given the highest expected level of resource competition. The negative control had higher C incorporation, but not higher cell abundances. We also examined an independent measure of growth through uptake of ^15^N-labeled leucine, a substrate commonly used to measure bacterial growth (43); we confirmed that the *Marinobacter* genome contains a leucine transporter (Supplementary Table S4). Outer well *Marinobacter* cells were more ^15^N enriched in the presence of *Devosia* compared to *Alcanivorax* (*P* = 0.016), and significantly less enriched than negative control (*P* < 0.001, Figure 4d). Interestingly, a contrasting enrichment between the inner and outer well *Marinobacter* was observed in the positive control (Supplementary Figure S6). This is likely due to a concentric design of the microplate where outer *Marinobacter* was able to access more ^15^N-labeled leucine from immersing media outside of the device. On the other hand, cell size of recipient strain was not statistically different (0.73-0.74 μm^2^, *P* = 0.84) except for inner *Alcanivorax* cells that were the largest (median 1.00 μm^2^, *P* < 0.001, Supplementary Figure S7). This indicates that in the absence of any interspecies interaction, *Marinobacter* cells may have been modestly more metabolically active but not enough to be reflected in increased growth rate or cell size over the time scale tested. Instead, the size differed by the well location where inner *Marinobacter* in the positive control were significantly larger than in the outer well by 1.36-fold (*P* < 0.001, Wilcoxon test).

Influencer abundances and incorporated algal C also varied significantly (Figure 4e), likely a result of strain specific C use efficiency. *Devosia* had higher cell abundances and C incorporation than *Alcanivorax*, but, despite this, it did not negatively influence abundances or C incorporation for recipient *Marinobacter*, suggesting that even in high numbers and activity, *Devosia* was a more cooperative influencer than *Alcanivorax*. Of course, this also indicates that *Alcanivorax* was an especially efficient competitor, since despite this low abundance and low net incorporation it was still able to negatively influence the C incorporation of the *Marinobacter* recipient (Fig.4b).

To compare algal C drawdown of both the influencer and recipient together, we combined cell counts, cell size, and single cell carbon assimilation measurements to calculate total C incorporation in the entire microplate (both influencer and recipient wells) over the incubation period, denoted as *C*_total_ (Supplementary Note S2). Among the three tested pairs of isolates, the combination of *Devosia* and *Marinobacter* showed the highest incorporation of algal C, whereas the masses by *Alcanivorax*-*Marinobacter* was the least and equivalent to our negative control with no bacterial cells in the inner wells (Figure 4f). The stark difference between the two influencers is somewhat unexpected, as the metabolomics data indicated that *Alcanivorax* consumed and produced a much more diverse set of metabolites from exudate than *Devosia*, which could have indicated a higher total net carbon assimilation from that exudate.

### Metabolite imaging highlights differences in metabolite production

Given our findings in the different recipient responses to our two influencers, we conducted a final experiment to complement our LC-MS/MS exometabolomics analysis. Here we compared total metabolite production on solid medium using MALDI imaging analysis of the three bacterial isolates grown as colonies on solid medium (*Alcanivorax* EA2, *Devosia* B7WZ, and *Marinobacter* 3-2) in isolation and adjacent to *P. tricornutum*. This revealed total metabolite production patterns that complement the LC-MS/MS exometabolomic analysis. Overall, 246 metabolites were detected from at least one bacterial isolate (Supplementary Table S5). When grown as a colony, *Alcanivorax* EA2 and *Marinobacter* 3-2 produced more metabolites (42 and 41 detected metabolites respectively) than *Devosia* B7WZ (13 detected metabolites). Of these, 30 of the metabolites were unique to *Alcanivorax* EA2 and 20 were unique to *Marinobacter*, but none were unique to *Devosia* B7WZ. We found that 48 metabolites were specifically produced during growth without algal contact, meaning that they were detected from a bacterium grown as a colony, but not from the same bacteria grown in co-culture with *P. tricornutum*. However, only nine metabolites were detected that were produced only by a co-culture and not by either the bacterial isolate or *P. tricornutum* itself grown in isolation.

## Discussion

Microbial community assembly is modulated by a web of dynamic and context-dependent interactions where metabolites, info-chemicals, and toxins mediate the relative fitness of species (44). For our system, invoking a solely bottom-up perspective – one that structures communities through resource competition alone – failed to reproduce observed bacterial growth patterns. To explore not only competitive interactions but also cooperative ones, we designed a series of experiments that isolated temporal and spatial interaction modes. We found that (1) metabolomic profiling and genome-based modeling predictions correlated on degrees of resource competition between bacterial pairs, (2) sequential interaction experiments supported predictions of relative competition between pairs with outliers identifying the cooperative interactions, and (3) porous microplate isotope tracing with live algae supported metabolic niche partitioning predictions with increased recipient algal carbon assimilation with the cooperative influencer relative to the competitive influencer.

Carbon assimilation was unexpectedly distinct between the two model influencers, highlighting the importance of activity-based measurements to examine interactions. We found expected differences in C flow to the *Marinobacter* recipient based on interaction predictions: *Marinobacter* took up less C with the competitive influencer (*Alcanivorax*) relative to the influencer with the complementary metabolic niche (*Devosia*). However, when expanded to examine C flow in the whole microplate, *Devosia* had much higher C assimilation and higher cell abundances than *Alcanivorax*, leading to an amplified effect on total C incorporation. While metabolic profiling did not predict this difference, respiration rates can differ by orders of magnitude for different marine taxa, and abundance can be decoupled from respiration rate (45), so this could explain differences in metabolic efficiencies between the influencers. For example, *Alcanivorax* can consume a diverse array of substrates, but this may come with a cost that leads to low bacterial growth efficiency (high respiration), since growth efficiency is dependent on both resource quality and availability (46). It is important to note the three influencer bacteria can co-exist at relatively high abundances in enrichment communities with *P. tricornutum* (30, 47), indicating that even the most competitive interactions do not lead to exclusion in these cases. If *Alcanivorax* has a low growth efficiency in a community, or excretes new compounds then made available to other bacteria, this could explain the co-existence. Since we did not measure respiration or DOC concentrations, we do not know the mechanism that led to more algal-derived carbon assimilated into bacterial biomass in the cooperative interaction compared to the competitive interaction.

Our approach yet does not elucidate a molecular mechanism for our representative cooperative interaction. Given that metabolic cross-feeding is thought to be the most prevalent of bacterial cooperative interactions (24-27), it is surprising that our data does not support cross-feeding as the mechanism for *Devosia*’s cooperative influence. LC-MS/MS metabolite profiling and MALDI are complimentary, capture distinct sets of metabolites (LC-MS/MS has higher resolution, lower detection limits, and detected lower m/z features, but MALDI can detect compounds lost in solid phase extraction), and the experiments represent growth on liquid and solid media respectively. Covering a relatively broad window of detection, both experiments showed that *Devosia* produced the fewest metabolite features of the bacterial isolates. Further, *Devosia* did not produce any identifiable unique metabolite features, or features that were consumed by *Marinobacter*. While it is possible that *Devosia* was producing a compound that we could not detect, such as small organic acids and sugars not captured by MALDI or solid phase extraction, an alternative cooperative mechanism involves degradation of inhibitory compounds. While existing literature suggests an ability of the *Devosia* genus to transform toxins or their intermediates such as deoxynivalenol (48), polycyclic aromatic hydrocarbons (49), or potentially hexachlorocyclohexane (50, 51), it remains to be seen whether *P. tricornutum* can produce these or similar compounds. Of note, *P. tricornutum* is known to synthesize a number of bactericidal compounds, including eicosapentaenoic acid (EPA) (52, 53), which is abundant and can constitute up to half of total fatty acid production (54). EPA is active against both Gram-positive and Gram-negative bacterial species, acting on membrane fluidity and subsequent loss of osmoregulation (55). The presence of one or more bactericidal compounds supplied by the host introduces several interesting caveats to how we might interpret our results. First, we are unaware of the sensitivity of each of our isolates to the compounds, or whether some strains might not only be resistant but also consumers or degraders. For example, the presence of the bactericidal compounds degrader as an influencer might promote the growth of a sensitive recipient, compared to a pairing with another sensitive influencer. Second, it is unclear whether bactericidal compounds are secreted by live *P. tricornutum* or associated with dead cell debris, as well as their uncharacterized diffusivities in the porous structure in our co-culture system. This might explain the discrepancies between our sequential media and porous microplate experiments. The mechanisms of host-antagonistic interactions should be better resolved to inform how predicted competitive and commensal relationships behave in different experimental designs.

The work also demonstrates how the proximity to the host is influential in bacterial physiology. Using the spatially designed microplate and strain *Marinobacter* sp. 3-2, we unexpectedly discovered that the bacteria accumulated more leucine when distant from the host as evidenced by its higher ^15^N enrichment. While leucine incorporation has been a measure for estimating protein synthesis (43, 56, 57), it is recently shown to accumulate inside a starving bacterial cell (58) by its metabolic cost due to high-energy phosphate bonds (59). Our co-culture exemplifies how two lifestyles by a single species can exist at the same time, one of which represented as holding algal C with a high growth rate and cell size (*Marinobacter* in the inner wells) and another as metabolically distinct through starving (*Marinobacter* in the outer wells). The finding contributes to our previous understanding of these bacteria under a microenvironment near an alga (the “phycosphere” (60)) where diffusing exudates are the primary source of carbon under a nutrient-scarce space.

Taken as a whole, our results demonstrate that starting by predicting metabolic niche partitioning and then identifying outliers is a feasible approach to reveal interactions occurring on top of resource competition. Further, our results indicate that these complementary and cooperative interactions can influence carbon flow, which has important implications for our predictions of C fate in algal-dominated systems.

## Materials and Methods

### Strains and culturing conditions

Axenic microalga *P. tricornutum* CCMP 2561 was chosen as the host model species throughout the study. Unless stated otherwise, the diatom was cultured in seawater media at 20 ºC exposed to a diurnal cycle of 12 h light/12 h dark with a light intensity of 200 μmol m^-2^ s^-1^. The media contains commercially available sea salts of 20 g l^-1^ added with f/2 inorganic nutrients without silicate (f/2-Si) or artificial seawater medium (ESAW). The culture was routinely transferred to a new medium every 2-3 weeks under a biosafety cabinet or laminar flow hood to reduce the chance of bacterial contamination. Contamination tests were carried out by streaking culture samples on marine broth agar every 2-3 weeks and by checking for presence of bacteria using epifluorescence microscopy every 6-12 months. Ten bacterial strains were previously isolated from an enrichment community from a *P. tricornutum* outdoor mesocosm (47, 61). Each isolate was either maintained in 10% Zobell Marine Broth with Instant Ocean Salts, transferred to new medium every 3-4 weeks, or by co-culturing with *P. tricornutum* in f/2-Si.

### Untargeted metabolomics of exometabolite production and consumption

#### Incubation and sample collection

To remove any residual organic carbon, 500 ml glass Erlenmeyer flasks and 20 ml glass test tubes were baked at 500 °C for 2 hours. For algal incubation to obtain spent medium, flasks were filled with 250 ml ESAW and inoculated with one week old *P. tricornutum* culture. Cultures were incubated with shaking (90 rpm, 22 °C; 12 h : 12 h, light : dark; 3,500 lux illumination) for one week. Cultures were then combined and filtered through a 0.22 μm pore-sized polyethersulfone (PES) membrane. Inorganic phosphate and nitrogen were replenished in the collected spent medium by adding 0.005 g l^-1^ NaH_2_PO_4_.H_2_O, 0.0375 g l^-1^ NaNO_3_ and 0.0236 g l^-1^ NH_4_Cl (30).For bacterial incubation, baked tubes were filled with the 10 ml spent medium and inoculated with 300 μl of bacterial isolate culture. Five replicates per isolate were inoculated and 12 additional tubes were left uninoculated as a baseline control. Cultures were incubated for two weeks under the same conditions as above. Cells were removed by filtering the cultures through the 0.22 μm pore-sized filters.

#### Exometabolomic Analysis

Metabolite samples were extracted and analyzed using liquid chromatography tandem mass spectrometry (LC-MS/MS) as previously described (30). Briefly, metabolites were extracted from filtrates using Bond Elut PPL columns (Agilent, Santa Clara, CA, USA) as previously described (30). Metabolite extracts were dried, resuspended in 150 μl methanol containing ^13^C- and ^15^N-labeled matrix control internal standards, filtered through a 0.2 μm pore-sized PES membrane filter and transferred to an autosampler vial. Detailed instrument information and LC-MS/MS conditions and parameters are given in Supplementary Table S6. LC-MS/MS data were analyzed with an untargeted approach. MZMine software (62) was used to identify features based on mass-to-charge ratio (m/z) values and retention times, analyzing positive and negative ionization mode data separately. MZMine analysis parameters are detailed in Supplementary Table S7. Global Natural Products Social Molecular Networking (GNPS) (63) was used to analyze the identified features, conduct molecular networking and putatively identify metabolites.

The LC-MS/MS features (162 features that were identified based on retention time and m/z values and were significantly above background) were further processed to compute an expected competitive interaction (ECI) strength between a recipient and an influencer bacterial strain (Equation 1) (5, 40, 41).

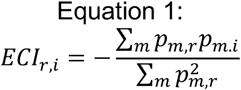

Here *ECI*_*r*,*i*_ is the effect on a recipient bacterial strain *r* from an influencer bacterial strain *i*, and *p*_*m*,*r*_, *p*_*m*,*i*_ are the proportions of metabolite *m* used by the influencer *i* or the recipient *r*, respectively. A proportion (e.g., *p*_*m*,*r*_ and *p*_*m*,*i*_) is equal to the ratio in LC-MS/MS signal intensities between isolate-inoculated and the uninoculated spent medium samples (5, 40, 41). Locations of identified LC-MS/MS features with peak heights, the GNPS results, and custom Python scripts for analyses are provided in Data, Materials, and Software Availability.

### Metabolic Modeling

#### Metabolic network reconstruction

Bacterial draft genome scale metabolic (GEM) models were reconstructed using CarveMe (64). The model was curated by an experimental determination to ensure it captured accurate phenotypes with a minimal set of growth supplements. To determine the supplements, 10 bacterial isolates were individually assayed for each of 190 sole carbon sources using 96-well Phenotype MicroArrays™ (Biolog Inc., Hayward, CA). Growth was monitored by absorbance at 600 nm for 96 hours in a plate reader and analyzed to classify whether a strain consumed a carbon source (Supplementary Note S3). For each isolate, a set of the growth-promoting carbon sources was used to generate a media database along with a base medium composition. The base medium contained macronutrients (NH_4_^+^, PO_4_^3-^, SO_4_^2-^), micronutrients (Co^2+^, Cu^2+^, Fe^2+^, Fe^3+^, Mn^2+^, MoO_4_^2-^, Ni^2+^, Zn^2+^), salts (Ca^2+^, Cl^-^, K^+^, Mg^2+^, Na^+^), and gasses (CO_2_, O_2_). The media database and protein fasta sequences for the enzymes in the metabolic models were passed as arguments to CarveMe.

#### Community metabolic modeling

Pairwise competitive interactions were quantified using metabolic resource overlap (*MRO*) (25). This score reflects the set of minimal nutrient requirements, denoted as *M*, shared between two species, *i, r* (Equation 2),

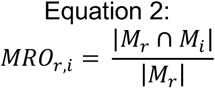

The MRO score reflects opportunities for direct competition, and does not consider the positive contributions of cross-feeding interactions. It relates most closely to the metabolite production and consumption experiments and is analogous to the ECI score.

### Pairwise bacterial isolate sequential spent media exchange

#### Incubation and sample collection

Eight hundred milliliters of *P. tricornutum* spent medium was prepared as described above. For the ‘influencer’ strain growth, eleven 125 ml flasks were prepared with 50 ml each of algal spent medium. Each flask was inoculated with 2 ml of a bacterial isolate culture. One flask was left uninoculated as a control. Flasks were incubated under the same conditions as above for two weeks. At the end of the ‘influencer’ incubation, cultures were filtered (0.2 μm pore size membrane) to remove cells. For ‘recipient’ strain growth, eleven 48-well plates (one per recipient strain plus control) were prepared with ‘influencer’ spent medium (3 wells per ‘influencer’, 750 μl per well). Each plate well was inoculated with 100 μl of a ‘recipient’ bacterial isolate culture (one isolate per plate) or no isolate (control) and incubated for two weeks. At the end of the ‘recipient’ incubation, samples (0.5 ml) were collected for flow cytometry from each well, fixed with glutaraldehyde (final concentration of 0.25%), and stored at - 80 °C.

#### Flow cytometry and analysis

Fixed samples were thawed and diluted in filter sterilized media (10% Zobell Marine Broth with Instant Ocean Salts) to achieve less than 10,000 counts per second. Diluted samples were aliquoted (250 μl) into a 96 well plate and stained with 2.5 μl of 100X SYBR Gold for 10-15 minutes in the dark. Cells were counted on an Attune benchtop flow cytometer as described previously (29). Blanks consisting of ultrapure water were run between each set of biological triplicates.

Using the bacterial cell counts, a sequential interaction (SI) representing the effect of an influencer on a recipient strain was calculated (Equation 3) (5).

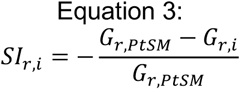

Here *SI*_*r*,*i*_ is the effect on a recipient *r* from an influencer *i, G*_*r*,*PtSM*_ is the growth (in cell counts) of recipient strain *r* on *P. tricornutum* spent medium, and *G*_*r*,*i*_ is the growth of recipient strain *r* on spent medium from influencer strain *i*. The difference between *G*_*r*,*PtSM*_ and *G*_*r*,*i*_ was normalized by the recipient growth so that SI strength scales to the ECI (Equation 1), i.e., -1 for a complete or 0 for no competitive impact.

### Porous microplate co-culture

#### Device preparation

A porous microplate was fabricated as previously described (31-33). Briefly, acrylic mold parts were laser cut from 1/8- and 1/4-inched acrylic sheet (Universal Laser Systems) and were attached by using acrylic (Weld-On) and epoxy (3M) adhesives. Using the acrylic mold, a polydimethylsiloxane (PDMS) mold was casted onto the acrylic and cured at 70 ºC for overnight. A prepolymer solution was prepared by mixing reagents as listed in Supplementary Table S8. The prepolymer was cast onto the PDMS mold, followed by placing a clean glass slide (75 × 50 mm^2^, VWR), and was polymerized by exposing to ultraviolet light with 365 nm wavelength for 15 min. Polymerized devices were detached from PDMS mold and stored in a glass jar containing 250 ml methanol. The methanol was replaced every day two times to remove remaining prepolymers. The copolymer devices were transferred to an autoclaved glass jar containing f/2-Si added with ^13^C sodium bicarbonate and ^15^N leucine to final concentrations of 2 mM; 10 nM, respectively, and were stored for 2-3 weeks before use.

#### Incubation and sample collection

As previously described (32), axenic *P. tricornutum* was acclimated to the copolymer by inoculating a stationary phase cells into a microplate. The cells were incubated for a week and were diluted four times using f/2-Si containing the isotopes. Diluted cells were inoculated into center well of a microplate at starting concentration of 8.4 × 10^6^ cells ml^-1^. For bacteria, each colony of *Alcanivorax* sp. EA2, *Devosia* sp. EAB7WZ and *Marinobacter* sp. 3-2 was inoculated to marine broth and grown overnight at 30 ºC, 250 rpm. Overnight cultures were washed twice with f/2-Si, sat overnight at room temperature and were diluted to OD600 ∼0.01 with isotope-containing f/2-Si. Diluted cells were inoculated into surrounding microplate wells. Porous microplates and the cells were immersed in f/2-Si with the isotopes and incubated. On days 5 and 14 post incubation, 35 μl bacteria and 300 μl *P. tricornutum* were collected from the microplate. Bacterial cells were subsampled and streaked on marine broth agar to confirm its presence and to test for cross-contamination. Remaining samples were fixed using formaldehyde with a final concentration of 2% v/v. Fixed cells were left at room temperature for a day and were stored at 4 ºC up to 8 weeks.

#### Flow cytometry

Forty microliters of fixed bacteria were added with 0.1 μl SYBR Green I nucleic acid stain and were allowed to sit for 0.5–1 h at room temperature without light exposure. To each well containing the stained cells, 2 μl flow cytometry counting beads and 158 μl 0.1 M TAPS buffer (pH 7.76) were added, bringing to a total volume of 200 μl. Flow cytometry was conducted on a BD FACS Canto II HTS and a BD FACS Diva software (Supplementary Table S9). Events were collected and clustered based on FITC-A and Alexa Flour 680-A gates. The count numbers were exported as csv files and analyzed using Microsoft Excel or R.

#### NanoSIMS imaging analysis

Twenty microliters were subsampled from each fixed sample with cells collected on day 14 post incubation. For each microplate and treatment, triplicates were pooled to bring to 60 μl in total and filtered on a small area of a 0.2 μm pore size polycarbonate membrane (Whatman Nuclepore, Cytiva, Marlborough, MA). Filters were rinsed, dried, the filtered areas cut and adhered to conductive carbon tape (Ted Pella, Redding, CA), gold coated, and analyzed by NanoSIMS as previously described (29, 47). Briefly, a ∼60 pA primary Cs^+^ beam was used to sputter a 33 x 33 μm raster for 2 cycles (131.1 s) prior to collecting 25 cycles over a 30 x 30 μm acquisition area with ∼2 pA Cs^+^ (∼150 μm diameter). The secondary ion mass spectrometer was tuned to ∼9500 mass resolving power (MRP, with aperture slit 2 and entrance slit 4), and automatic secondary ion beam centering was performed at each new location based on ^12^C^14^N^-^. Secondary ion images were collected for masses ^12^C^12^C^-, 12^C^13^C^-, 12^C^14^N^-, 12^C^15^N^-^, and ^32^S^-^ on individual electron multipliers, as well as secondary electrons (SE). All nanoSIMS images were processed using L’Image software to correct for dead time and image shift across cycles, create ^13^C/^12^C and ^15^N/^14^N ratio images, and draw regions of interest (ROIs) to measure bacterial ^13^C and ^15^N incorporation. We calculated isotope ratios (^12^C^13^C^-^ / ^12^C^12^C^-^ × 0.5 = ^13^C / ^12^C) of each bacterial cell (34). Based on the algal ^13^C labeling, which is the source of C for the bacteria, the percent of bacterial carbon derived from the alga was calculated (net carbon incorporation, C_net_) (29, 35, 42). Since the isotope composition of algal exudate could differ from the measured ^13^C enrichment of the algal cells, herein C_net_ is considered as a comparative estimate of bacterial incorporation. We also note that algal cells were similarly abundant across treatments (Supplementary Figure S8), which should lead to similar levels of isotope enrichment. A total incorporation of algal C mass by bacterial isolates was defined and calculated (Supplementary Note S2), which is by combining the single cell C_net_ and their abundances across the microplate culture wells.

### Matrix assisted laser desorption/ionization (MALDI) imaging of algal and bacterial metabolites

#### Sample preparation

Axenic *P. tricornutum*, three bacteria only (*Devosia, Alcanivorax*, and *Marinobacter*) and three co-cultures (*P. tricornutum* with *Devosia*, with *Alcanivorax*, and with *Marinobacter)* were incubated in liquid f/2 media for one week. Ten microliters of each culture were spotted onto f/2 agar, allowed to dry for 10 minutes. The plates were wrapped in parafilm and incubated under a 14 h:10 h light/dark cycle at 22 °C for 10 days.

#### MALDI mass spectrometry imaging (MSI) and analysis

Areas with interaction colonies were excised from agar, placed onto a double-sided adhesive copper tape (3-6-1182; 3M) adhered to indium tin oxide-coated glass slides (Bruker Daltonics, Billerica, MA), and dried at room temperature overnight prior to MALDI matrix application and MALDI MSI analysis. MALDI matrix (DHB: 40 mg/mL in 70% MeOH) was applied to the ITO-coated glass slide using an HTX TM-Sprayer (HTX Technologies). Imaging was performed on a 15 tesla (T) MALDI-Fourier transform ion cyclotron resonance-mass spectrometer (FTICR-MS; SolariX, Bruker Daltonics) equipped with SmartBeam II laser source (355 nm, 2 kHz). Spraying, imaging, and ion collection conditions are detailed in Supplementary Table S10. MS imaging data ware acquired using FlexImaging (v4.1, Bruker Daltonics), images were processed and visualized via SCiLS Lab (v2024a Premium 3D, Bruker Daltonics) from the profile datasets. The centroided dataset was automatically annotated within METASPACE (parameters given in e) and its list of features was imported back to the SCiLS after RMS normalization. Colony borders were manually drawn based on flatbed scanner image prior to MALDI imaging analysis. Features for each colony were filtered based on ion mobility’s intensity being greater than 10,000 above 3 times the intensity of blank agar.

## Data, Materials, and Software Availability

LC-MS/MS, NanoSIMS and flow cytometry data including custom scripts for visualization are available at GitHub repository (https://github.com/vbrisson/phycosphere_bacteria-bacteria_interactions). Information on the GNPS analysis and results are available at https://gnps.ucsd.edu/ProteoSAFe/status.jsp?task=94d9cced1af245bcbf6216e66fc91720 for positive mode data and https://gnps.ucsd.edu/ProteoSAFe/status.jsp?task=7a90fe1862d945ec84f8d48647f00fdf for negative mode data. Codes for draft genome scale metabolic models are provided at GitHub repository PheArrMe (https://github.com/jrcasey/PheArrMe). MALDI mass spectrometry imaging data available at https://metaspace2020.eu/project/algae-bacteria_MSI. All other data are provided in Supplementary Materials.

## Acknowledgments

This research was funded by U.S. Department of Energy (DOE) Office of Biological and Environmental Research, SCW1039, under the microBiospheres bioenergy Science Focus Area. The work at LLNL was performed under DOE auspices of the by Lawrence Livermore National Laboratory under Contract DE-AC52-07NA27344. MALDI-MSI data was collected as part of project award 50220 under the Facilities Integrating Collaborations for User Science (FICUS) program and used resources at the Environmental Molecular Sciences Laboratory (EMSL), which is a DOE Office of Science User Facility operated under contract DE-AC05-76RL01830. Bacterial genomes were sequenced through the Joint Genome Institute (JGI) Community Sequencing Program award number 939. Work conducted by the JGI, a DOE Office of Science User Facility, is operated under Contract No. DE-AC02-05CH11231. H.K. was partly supported by the Kwanjeong Educational Foundation. We thank the Koch Institute’s Robert A. Swanson (1969) Biotechnology Center, specifically Glenn Paradis and Michele Griffin at The MIT Flow Cytometry Core.

