## Supplementary Information for "Spatially structured competition and cooperation alters algal carbon flow to bacteria"

1 Department of Mechanical Engineering, Massachusetts Institute of Technology, Cambridge, MA, USA; 2 Physical and Life Sciences Directorate, Lawrence Livermore National Laboratory, Livermore, CA, USA; 3 Environmental Genomics and Systems Biology Division, Lawrence Berkeley National Laboratory, Berkeley, CA, USA; 4 The DOE Joint Genome Institute, Lawrence Berkeley National Laboratory, Berkeley, CA, USA; 5 The Environmental Molecular Sciences Laboratory, Pacific Northwest National Laboratory, Richland, WA, USA.

* Equal contributions.

† To whom correspondence may be addressed.

List of Supplementary Materials

Supplementary Figure S1. LC-MS/MS metabolomics profiles for bacterial isolated grown on spent medium from *P. tricornutum*.

Supplementary Figure S2. Relationship between metabolic resource overlap (MRO) and estimated competitive interactions (ECI).

Supplementary Figure S3. Design of porous microplate for algal-bacterial co-culture.

Supplementary Figure S4. Carbon net incorporation of bacteria co-cultured with *P. tricornutum* in porous microplate.

Supplementary Figure S5. Abundances of bacteria co-cultured with *P. tricornutum* in porous microplate.

Supplementary Figure S6. Enrichment of $^{15}N$ by *Marinobacter* in porous microplate.

Supplementary Figure S7. Cell area of *Marinobacter* co-cultured with *P. tricornutum* in porous microplate.

Supplementary Figure S8. Abundance of *P. tricornutum* incubated in porous microplate.

Supplementary Table S1. LC-MS/MS feature peak heights.

Supplementary Table S2. Putative identifications for metabolite features based on spectral matches in GNPS.

Supplementary Table S3. Flow cytometry counts from the spent medium exchange experiment used to calculate the sequential interaction strength (SI).

Supplementary Table S4. Top five matches from sequence alignments between *Marinobacter* genome and bacterial leucine transporter gene.

Supplementary Table S5. Feature detection with matrix assisted laser desorption/ionization (MALDI) imaging.

Supplementary Table S6. Instrument information and LC-MS/MS parameters for metabolomic analysis.

Supplementary Table S7. MZMine analysis parameters for feature identification.

Supplementary Table S8. Reagents and their composition for copolymer poly(2-hydroxethyl methacrylate-co-ethylene glycol dimethacrylate) (HEMA-EDMA).

Supplementary Table S9. Parameters for operating flow cytometry and software.

Supplementary Table S10. Parameters and conditions for operating MALDI mass spectrometry imaging (MSI) and analysis.

Supplementary Note S1. Design of porous microplate and estimation of algal exometabolites diffusion.

Supplementary Note S2. Derivation of total algal carbon mass incorporation by bacteria in porous microplate.

Supplementary Note S3. Determination of minimal bacterial growth supplements for metabolic model reconstruction.

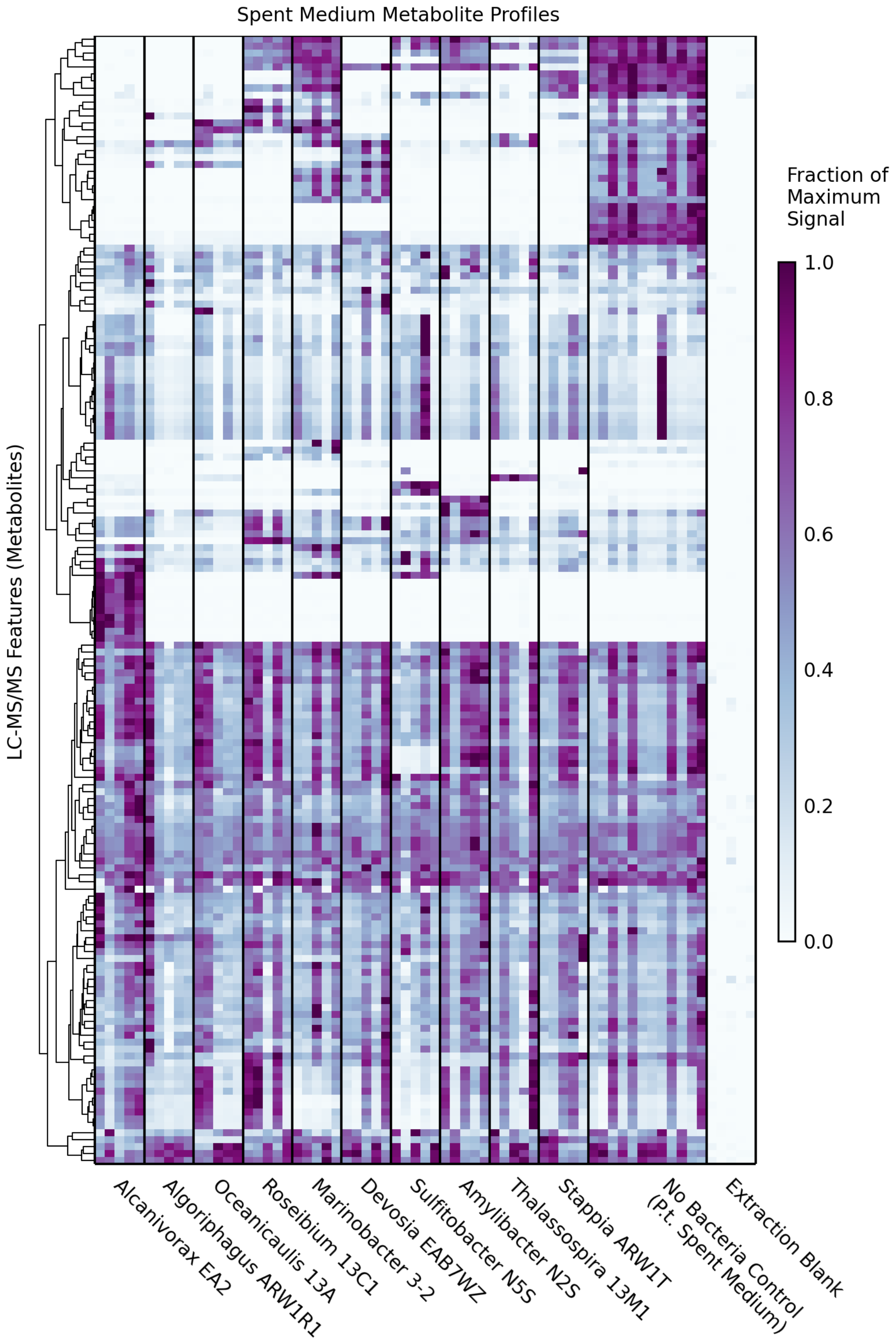


**Supplementary Figure S1.** LC-MS/MS metabolomics profiles for bacterial isolated grown on spent medium from *P. tricornutum*. Each row represents one LC-MS/MS feature (*m/z* and retention time), and each column represents one sample. Samples are grouped by the bacterial isolate, with all 5 replicate samples shown. Rows are grouped based on hierarchical clustering, as indicated in the dendrogram on the left.

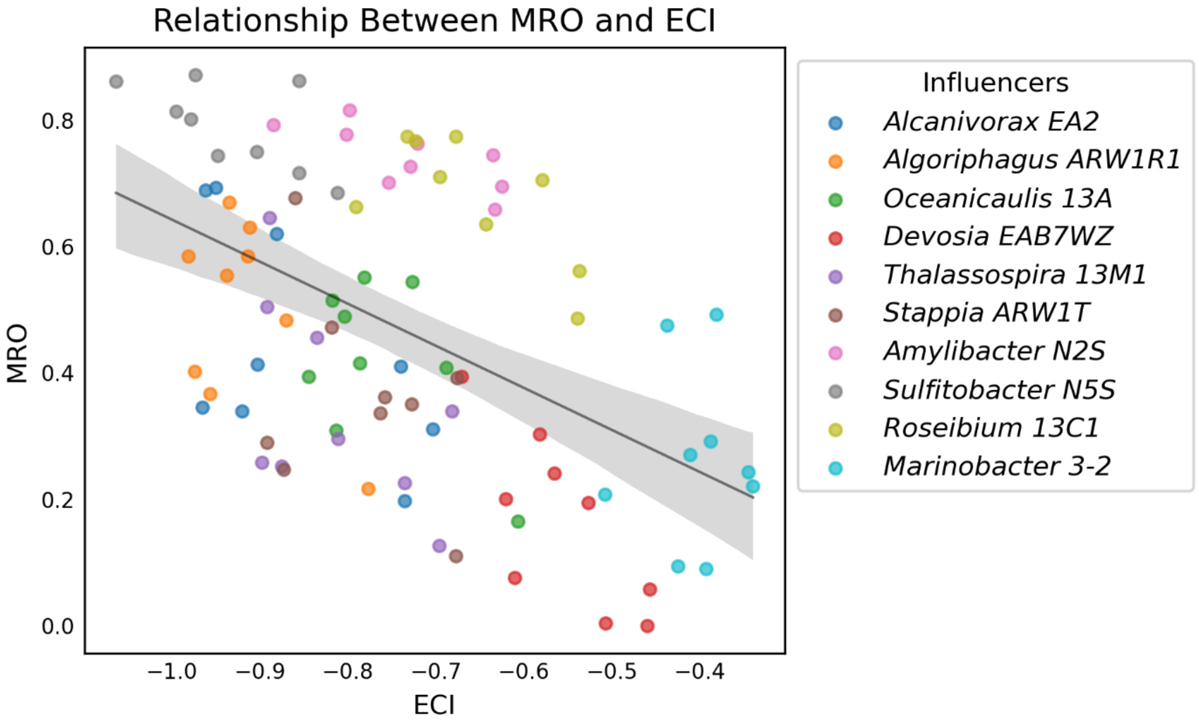


**Supplementary Figure S2.** Relationship between metabolic resource overlap (MRO) and estimated competitive interactions (ECI). Each point represents one influencer-recipient pair, and the color is based on the influencer strain. The diagonal line shows the Pearson correlation ($R^2 = 0.25$, $P = 6 \times 10^{-7}$), and the grey area indicates the 95% confidence interval.

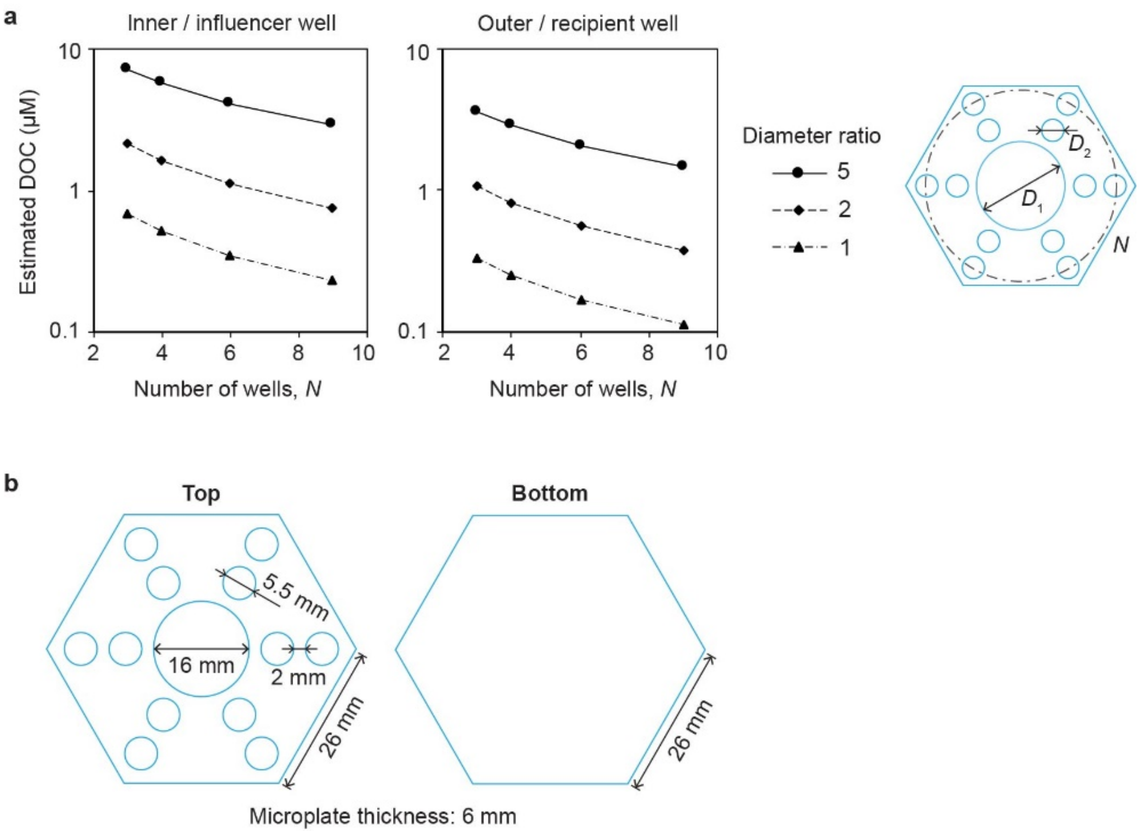


**Supplementary Figure S3.** Design of porous microplate for algal-bacterial co-culture. (a) Estimation of log-scaled DOC concentration diffused to surrounding wells, parametrized by a ratio between center and surrounding wells and the number of surrounding wells for each distance level (inner, left; outer, right). Concentrations are calculated based on abundance of *P. tricornutum* incubated in the porous microplate for twenty days (see Supplementary Note S1 for details). (b) Final dimension of porous microplates used in this study.

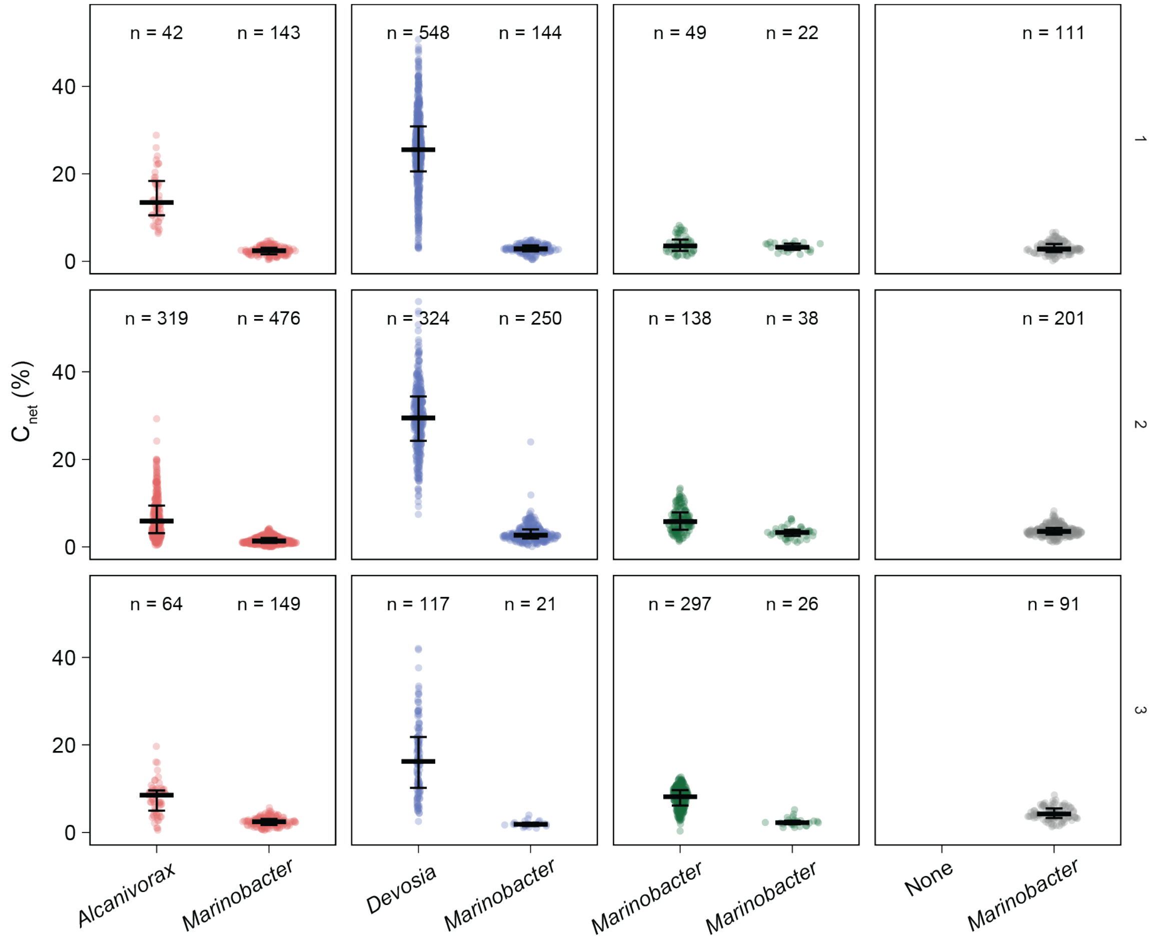


**Supplementary Figure S4.** Net carbon incorporation ($C_{net}$) of bacteria co-cultured with *P. tricornutum* in porous microplate measured using single-cell isotope tracing and NanoSIMS. Each facet displays a pair of co-cultured isolates as outlined in Table 2 of the main text with replicating microplates for each row. Black middle line and error bar indicate median and interquartile ranges, respectively. Number of single cells (n) analyzed for each treatment or location is displayed.

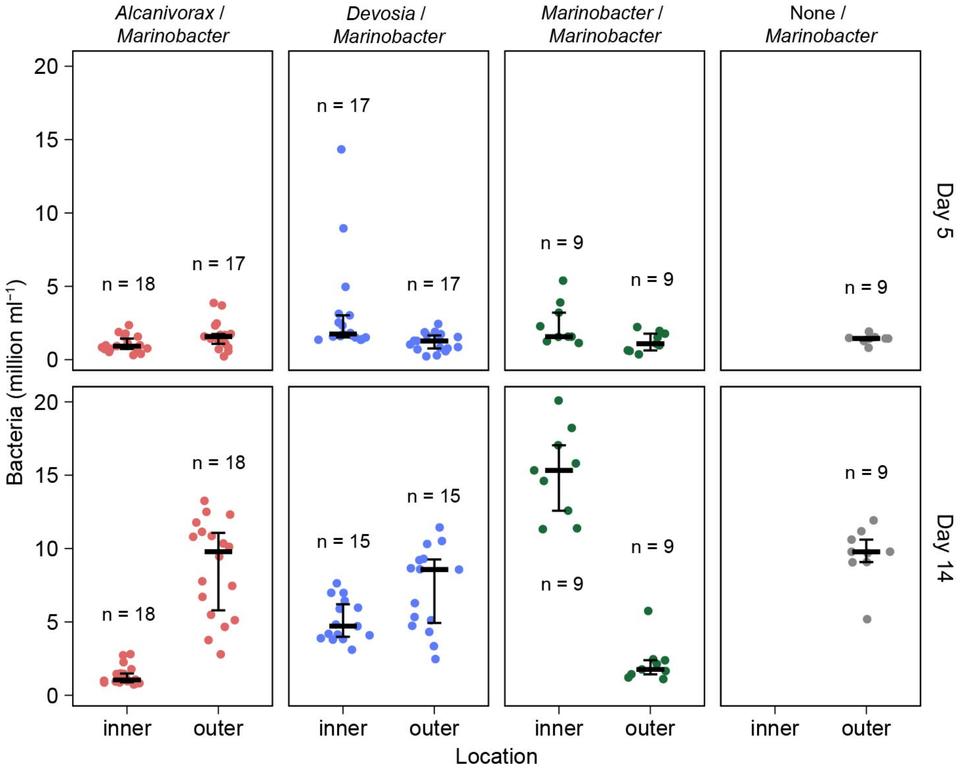


**Supplementary Figure S5.** Abundances of bacteria co-cultured with *P. tricornutum* in porous microplate. Each point represents measurement using flow cytometry of samples from each well of the microplate on days 5 and 14 of incubation (top and bottom row). Black lines indicate median (thick) and interquartile ranges (error bar). Number of single cells (n) analyzed for each treatment or location is displayed.

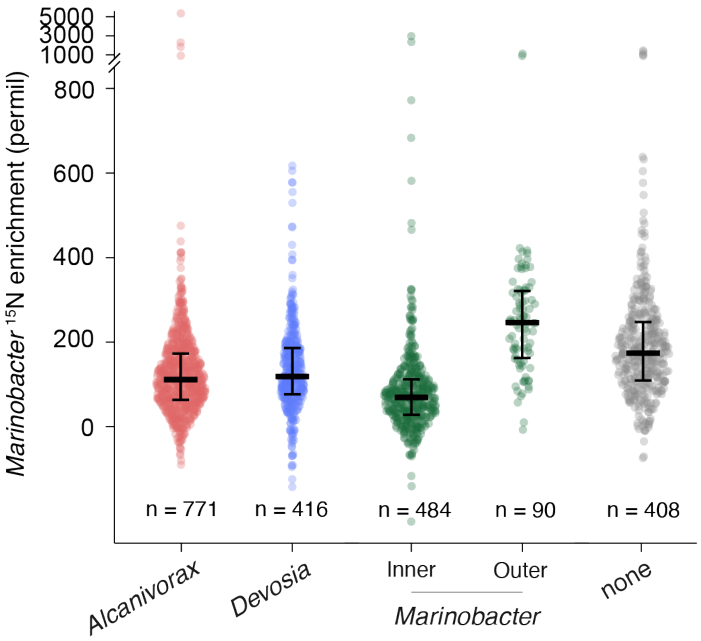


**Supplementary Figure S6.** Enrichment of $^{15}N$ by Marinobacter sp. 3-2 co-cultured with influencer strains (x axis) in porous microplate. The enrichment was measured using single-cell isotope tracing and NanoSIMS. Black middle line and error bar indicate median and interquartile ranges, respectively. Number of single cells (n) analyzed for each treatment or location is displayed.

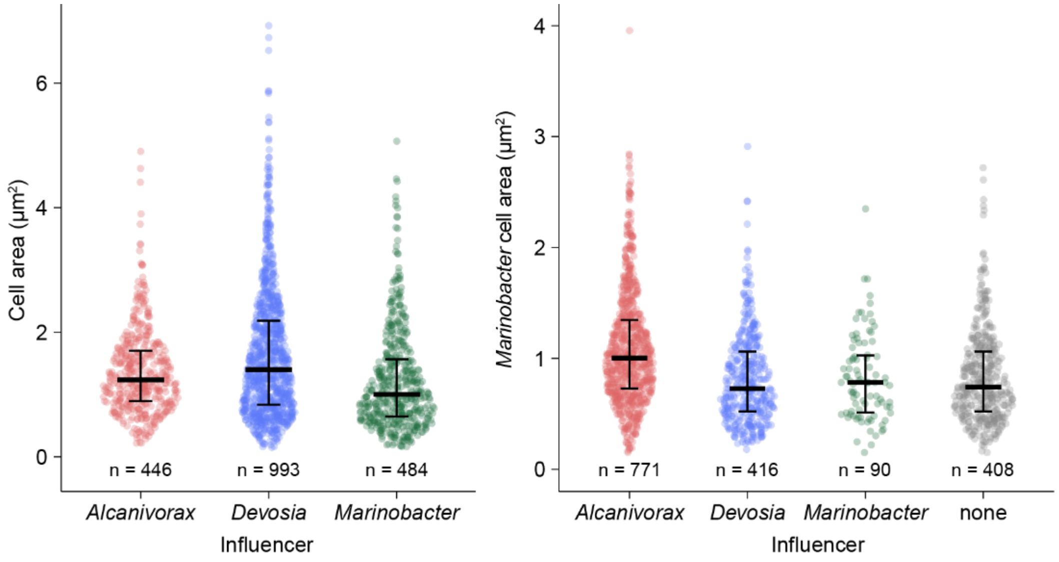


**Supplementary Figure S7.** Individual cell area of *Marinobacter* co-cultured with *P. tricornutum* in porous microplate. Black lines indicate median (thick) and interquartile ranges (error bar). Each point represents measurement of single cell area region of interest (ROI) from nanoSIMS images. Number of data is displayed for each treatment or location.

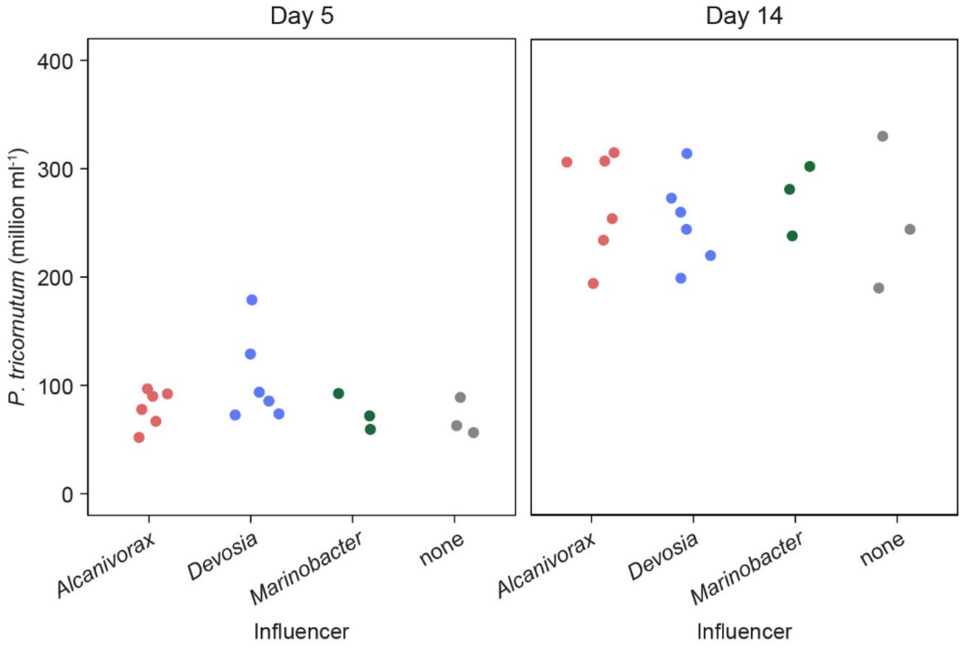


**Supplementary Figure S8.** Abundance of *P. tricornutum* incubated in porous microplate. *P. tricornutum* was incubated for day 5 and 14 and co-cultured with different bacterial influencer strain and recipient strain *Marinobacter*, located in inner and outer wells respectively.

**Supplementary Note S1. Design of porous microplate and estimation of algal exometabolites diffusion.**

Previous work employing the co-culture porous microplate have demonstrated that its building material, i.e., porous copolymer, enables microbial cells to communicate by exchanging their metabolites for a period of growth (~ weeks), while preventing a direct contact between adjacent cultures. We further characterized the copolymer by measuring its biomolecular diffusivity [1], tortuosity, porosity and by visualizing its structure [2]. Based on its characterization a numerical model was established estimating a concentration of algal photosynthates diffused to surrounding wells during incubation [3]. In specific, concentration of dissolved organic carbon (DOC) exuded by alga *P. tricornutum* was estimated based on its growth experimentally measured in the microplate, and Fick's law of diffusion was used to calculate the DOC concentration in adjacent or surrounding wells.

In this work, the model is parametrized by changing 1) the number of surrounding wells for each distance level from the center and 2) the size ratio between center and surrounding well (Figure 3a). The objective of this parameter study was to find an optimal configuration of a porous microplate for testing our hypothesis on sequential, bacterial uptake of algal exometabolites. Therefore, by designing the microplate we sought to maximize the amount of exometabolites reaching the bacteria, and at the same time, retain sufficient replicates for reliability. Numerical results show the concentration increased by decreasing number of surrounding wells and increasing well size ratio. The results also show that the change in the surrounding well number had less impact than the size ratio in the concentration. These findings allowed us to configure the microplate design with six surrounding wells and the highest size ratio (e.g., 3, Supplementary Figure S4b).

**Supplementary Note S2. Derivation of total algal carbon mass incorporation by bacteria in porous microplate.**

We derive the total algal carbon mass incorporated by bacterial isolates in a porous microplate. Denoted as $C_{\text{total}}$, the total mass incorporation is a summation by the two isolates co-cultured in the microplate, $C_i$ and $C_r$, with each subscript respectively denoting the influencer ($i$) and the recipient ($r$). Each incorporation is defined as, for example,

$$C_i(i,r) = \mathrm{C}_{\text{net},\, i}(i,r) \times N_i(i,r) \times V \times M_i,$$

where $C_i(i,r)$ is the incorporation by influencer $i$ with the presence of recipient $r$, $\mathrm{C}_{\text{net},\, i}(i, r)$ the influencer's average single cell $\mathrm{C}_{\text{net}}$, $N_i(i, r)$ the number of the cells per volume, $V$ the culture volume, $M_i$ the mass of a single cell. The relation allows to combine single-cell algal carbon uptake and bacterial abundance in the microplate co-culture. Numerical values for calculating $C_{\text{total}}$ for each bacterial pair are detailed in Data, Materials, and Software Availability of main text.

**Supplementary Note S3. Determination of minimal bacterial growth supplements for metabolic model reconstruction.**

Carbon sources as growth supplements were determined by using the 96-well formatted Phenotype MicroArrays™ (Biolog Inc., Hayward, CA). In brief, each of 10 bacterial isolates was inoculated to a microplate containing 190 carbon substrates in each well following manufacturer's protocol (95 substrate and one blank control per microplate). All wells contained 3% sodium chloride dissolved in deionized water. Bacterial growth was measured using plate reader for 96 hours. Absorbance time-series profiles were variable across isolates in both shape and range, and the distribution of absorbance ranges (both linear and log transformed) was not bimodal, making a filtering threshold arbitrary. In the absence of a more sophisticated approach to filtering, a conservative classifier was implemented, wherein growth was assessed if the time-series profile met the following criteria: (a) $t_{a_i}^{max} > t_{a_i}^{min}$ and $\max a_i(t) - \min a_i(t) > f(\max a_{neg}(t) - \min a_{neg}(t))$$t_{max(ai)} > t_{min(ai)}$ and max($a_i$(t)) – min($a_i$(t)) > f(max($a_{neg}$(t)) – min($a_{neg}$(t))), where $a_i$(t) is the absolute absorbance value at time t for the $i^{th}$ carbon source, $a_{neg}$(t) is the absolute absorbance at time t for the negative control, and f is a scaling factor. A visual inspection of the data was used to identify an appropriate scaling factor (f).
